# Dissection of the cellular function of the ZBED6 transcription factor in mouse myoblast cells using gene editing, RNAseq and proteomics

**DOI:** 10.1101/540914

**Authors:** Shady Younis, Rakan Naboulsi, Xuan Wang, Xiaofang Cao, Mårten Larsson, Ernest Sargsyan, Peter Bergsten, Nils Welsh, Leif Andersson

## Abstract

The transcription factor ZBED6 acts as a repressor of *Igf2* and affects directly or indirectly the transcriptional regulation of thousands of genes. Here, we use gene editing in mouse C2C12 myoblasts and show that ZBED6 regulates *Igf2* exclusively through its binding site 5′-GGCTCG-3′ in intron 1 of *Igf2*. Deletion of this motif (*Igf2*^*ΔGGCT*^) or complete ablation of *Zbed6* leads to ~20-fold up-regulation of IGF2 protein. Quantitative proteomics revealed an activation of Ras signaling pathway in both *Zbed6^-/-^* and *Igf2*^*ΔGGCT*^ myoblasts, and a significant enrichment of mitochondrial membrane proteins among proteins showing altered expression in *Zbed6^-/-^* myoblasts. Both *Zbed6^-/-^* and *Igf2*^*ΔGGCT*^ myoblasts showed a faster growth rate and developed myotube hypertrophy. These cells exhibited an increased O_2_ consumption rate, due to IGF2 up-regulation. Transcriptome analysis revealed ~30% overlap between differentially expressed genes in *Zbed6^-/-^* and *Igf2*^*ΔGGCT*^ myotubes, with an enrichment of up-regulated genes involved in muscle development. In contrast, ZBED6-overexpression in myoblasts led to cell apoptosis, cell cycle arrest, reduced mitochondrial activities and ceased myoblast differentiation. The similarities in growth and differentiation phenotypes observed in *Zbed6^-/-^* and *Igf2*^*ΔGGCT*^ myoblasts demonstrates that ZBED6 affects mitochondrial activity and myogenesis largely through its regulation of IGF2 expression. This study suggests that the interaction between ZBED6-*Igf2* may be a therapeutic target for human diseases where anabolism is impaired.

## INTRODUCTION

The ZBED6 transcription factor is unique to placental mammals and has evolved from a domesticated DNA transposon located in the first intron of *ZC3H11A*, a zinc-finger protein with RNA-binding capacity (Markljung et al., 2009; Younis et al., 2018a). ZBED6 was identified as a repressor of insulin-like growth factor 2 (*IGF2)* expression following the identification of a mutation in *IGF2* intron 3 in domestic pigs (Van Laere et al., 2003; Markljung et al., 2009). This mutation disrupts a ZBED6 binding site and leads to a 3-fold increase in *IGF2* mRNA expression in pig skeletal muscle, which in turn results in increased muscle mass and reduced subcutaneous fat deposition. We have previously reported that ZBED6 has thousands of putative binding sites in human and mouse genomes, with a strong enrichment in the vicinity of transcription start sites (TSS) of genes involved in development and transcriptional regulation (Markljung et al., 2009; Jiang et al., 2014; Akhtar Ali et al., 2015; Wang et al., 2018). However, it is still unknown which of these genes, besides *Igf2*, are true functional targets of ZBED6. Silencing of *Zbed6* expression in mouse C2C12 myoblasts using small interfering RNA (siRNA) resulted in differential expression of about 700 genes, including a 3-fold up-regulation of *Igf2* mRNA (Jiang et al., 2014). IGF2 is an essential growth factor in skeletal muscle development and has a role in the initiation of myoblast differentiation (Florini et al., 1991). Recently, we have developed *Zbed6^-/-^* and *Igf2* knock-in mice, the latter carrying the pig mutation at the ZBED6 binding site. These mice exhibited increased body weight and skeletal muscle growth (Younis et al., 2018b). However, the molecular mechanism how ZBED6 affects muscle growth has not been fully investigated. Particularly, the interaction between ZBED6 and *Igf2* during myogenesis, and to which extent phenotypic changes associated with altered ZBED6 expression is mediated through its interaction with the *IGF2* locus has hitherto not been studied.

In recent years, the genome editing technique based on the microbial Clustered Regularly Interspaced Short Palindromic Repeats (CRISPR) and CRISPR associated protein 9 (Cas9) nucleases, has become the most efficient method to knock-out a gene of interest or manipulate a specific site in mammalian cells (Jinek et al., 2012; Ran et al., 2013). We employed this technology to explore the significance of ZBED6-*Igf2* interaction in the mouse myoblast C2C12 cell line that has the ability to differentiate and form myotubes (Yaffe and Saxel, 1977). In the present study, we generated two models of engineered C2C12 cells, a *Zbed6* knock-out and a deletion of the ZBED6 binding site in an *Igf2* intron. The genetically modified C2C12 cells were induced to differentiate, followed by whole transcriptome analysis, mass spectrometry (MS)-based quantitative proteomics and detailed functional characterizations of myoblast proliferation and myotube formation.

## RESULTS

### Efficient deletion of *Zbed6* and its binding site in *Igf2*

In order to explore to which extent phenotypic changes associated with altered expression of ZBED6 is due to its interaction with the *Igf2* locus, we established two genetically engineered C2C12 cell lines, one with complete *Zbed6* inactivation and the other with a deletion of the ZBED6 binding site in the first intron of the *Igf2* gene (chr7:142,664,244-142,664,249, mmu10). The CRISPR/Cas9 method was used to delete almost the entire coding sequence (2.5 kb out of 2.9 kb) of *Zbed6* in C2C12 cells (Figure 1A). We genotyped 150 clones using multiplex PCR (Figure 1B) and found that 10% of the clones were untargeted, while 90% were targeted with at least one guideRNA. Of these clones, 22% showed a frameshift in one allele, 74% showed a 2.5 kb deletion of ZBED6 in single allele and 4% of the clones showed a 2.5 kb deletion in both alleles.

**Figure 1.**
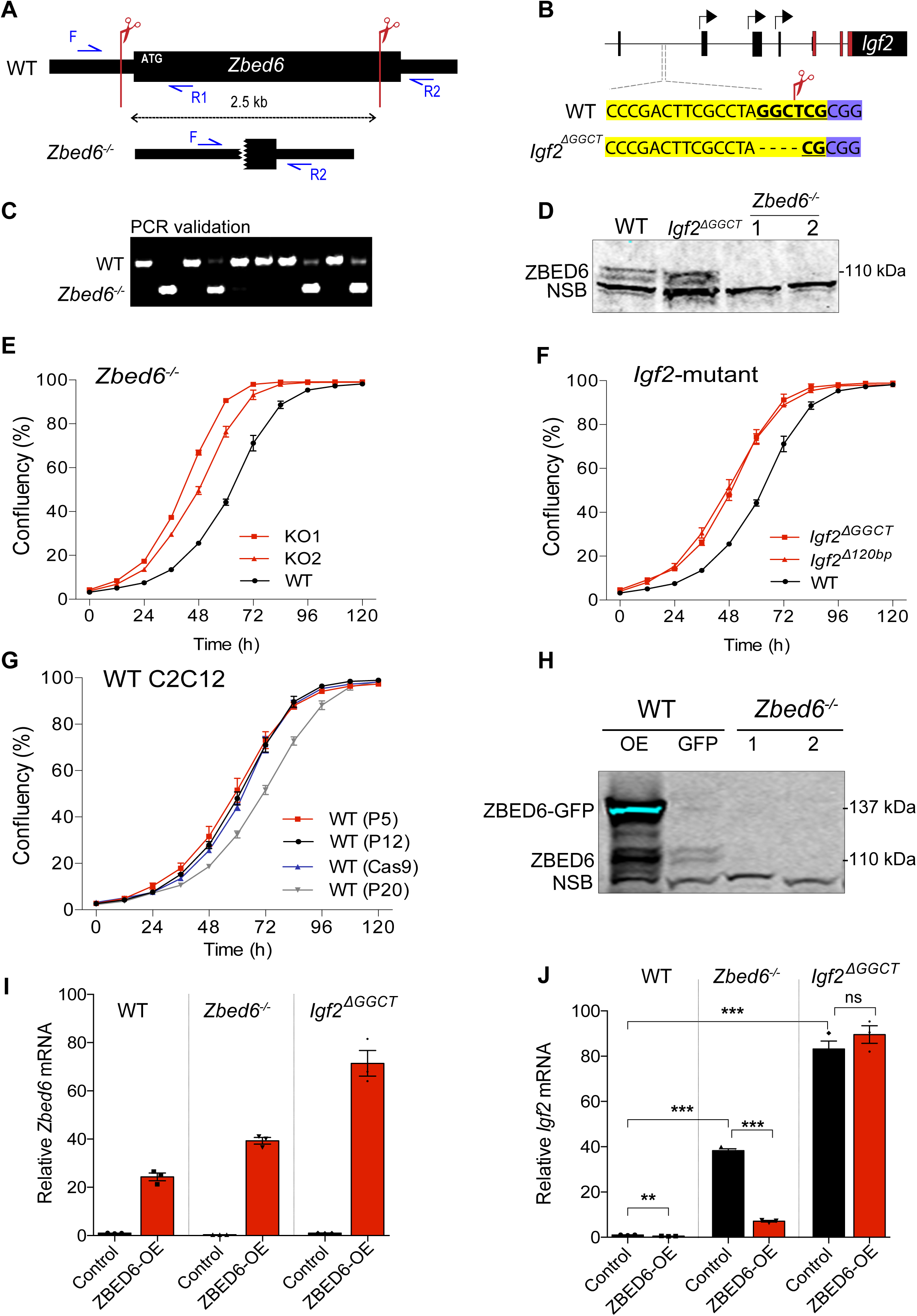
Knock-out of *Zbed6* or its binding site in *Igf2* alter the growth of myoblasts. (A) Schematic description of Zbed6 targeting using CRISPR/Cas9. Red scissors indicate the targeted sites of *Zbed6* using two guide RNAs. Blue arrows indicate the location of the PCR primers that were used for genotyping of the KO clones. (B) PCR screening of *Zbed6* KO clones. (C) Schematic description of the targeted ZBED6 binding sequences in *Igf2* (bold). The scissor indicates the cleavage site using specific gRNA sequences (yellow) adjacent to the PAM sequences (blue). Black arrows indicate *Igf2* promoters, red boxes are the coding sequences of Igf2. (D) Immunoblot validation of *Zbed6-/-* clones, the *Igf2*^*ΔGGCT*^ clone and WT cells, NSB: non-specific band. (E) Real-time measurements of cell growth (mean±SEM) of WT C2C12 cells (black) and *Zbed6^-/-^* clones (red) (n=3). (F) Cell growth of two *Igf2*-mutant clones (red) and WT cells (black). (G) Cell growth measurement of WT C2C12 cells at different passages (P5, P12 and P20) and WT cells transfected with Cas9 reagents without gRNA (WT Cas9). (H) Immunoblot validation of ZBED6-GFP overexpression in C2C12 cells. (I) Quantitative PCR analysis of *Zbed6* mRNA expression and after transient expression of GFP (Control) or ZBED6-GFP (ZBED6-OE) constructs in myoblasts. (J) Quantitative PCR analysis of *Igf2* mRNA expression after transient expression of ZBED6-GFP in WT cells*, Zbed6^-/-^* and *Igf2*^*ΔGGCT*^ cells. Graph shows the fold-changes (mean ± SEM) compared to WT control cells. ns=non significant, **P<0.01, ***P<0.001, Student’s t-test.

The ZBED6 binding motif 5′-GGCTCG-3′ in *Igf2* intron 1 was targeted using a guideRNA that cleaves between the C and T nucleotides (Figure 1C). PCR screening of C2C12 clones revealed several clones with deletions at this site, ranging in size from 4 to 120 nucleotides. One of the clones showed a four-nucleotide deletion (GGCT) of the binding motif in both alleles. We named this clone *Igf2*^*ΔGGCT*^ and used it for all downstream analysis together with the *Zbed6^-/-^* clones; a second clone (*Igf2*^*Δ120bp*^) was used to confirm the effect on cell growth (see below). The expression of the ZBED6 protein in the *Zbed6^-/-^* and *Igf2*^*ΔGGCT*^ clones was evaluated by immunoblot analysis, which revealed complete ablation of ZBED6 expression in *Zbed6^-/-^* clones and normal expression in *Igf2*^*ΔGGCT*^ cells in comparison to wild-type (WT) C2C12 cells (Figure 1D).

### Both *Zbed6^-/-^* and *Igf2*^*ΔGGCT*^ myoblasts exhibit faster cell growth and massive up-regulation of *Igf2* expression

The real-time measurement of growth rate showed that both *Zbed6^-/-^* and *Igf2*^*ΔGGCT*^/*Igf2*^*Δ120bp*^ clones grow faster than the WT (Cas9) cells (Figures 1E and 1F); the WT (Cas9) cells here refer to WT cells treated with CRISPR/Cas9 reagents without guideRNA and kept at similar selection conditions as the targeted cells. In order to explore the effect of this condition on cell growth, we measured the growth rate of WT (Cas9) versus the WT C2C12 cells at early (P5), middle (P12) and late (P20) cell passages. The WT (Cas9) had similar growth rate as the WT C2C12 cells at early and middle passage, while the late passage C2C12 cells grew slower (Figure 1G).

Expression analysis using quantitative reverse transcriptase PCR (RT-qPCR) revealed a more than 30-fold up-regulation of *Igf2* mRNA when ZBED6 was deleted or its binding site in *Igf2* was disrupted (Figure 1J). To verify that the increased expression of *Igf2* in *Zbed6^-/-^* and *Igf2*^*ΔGGCT*^ cells indeed was caused by the loss of ZBED6 or its binding site, we generated and validated an expression construct that produces a ZBED6-GFP fusion protein (Figure 1H). We reintroduced ZBED6 into the *Zbed6^-/-^* and *Igf2*^*ΔGGCT*^ clones by transient overexpression of ZBED6-GFP (ZBED6-OE). The GFP construct was used as control. qPCR analysis confirmed an efficient overexpression of *Zbed6* in WT, *Zbed6^-/-^* and *Igf2*^*ΔGGCT*^ cells (Figure 1I). The overexpression of *Zbed6* in WT cells resulted in a significant 60% down-regulation of *Igf2* mRNA. Interestingly, reintroduction of ZBED6-GFP in *Zbed6^-/-^* clones significantly down-regulated the elevated expression of *Igf2*, while no changes were observed when ZBED6-GFP was over-expressed in *Igf2*^*ΔGGCT*^ clones (Figure 1J). These results imply that ZBED6 represses *Igf2* expression exclusively through interaction with its binding site in *Igf2* intron 1.

### *Zbed6^-/-^* and *Igf2*^*ΔGGCT*^ myoblasts develop myotube hypertrophy with improved contractile properties after differentiation

The differentiation profile of *Zbed6^-/-^* and *Igf2*^*ΔGGCT*^ myoblasts was assessed by immunofluorescence staining of myotubes using myogenin and myosin heavy chain (MyHC) antibodies (Figure 2A). The differentiation of *Zbed6^-/-^* and *Igf2*^*ΔGGCT*^ myoblasts resulted in formation of hypertrophic myotubes (Figure 2A), with significant increase in the differentiation index in *Zbed6^-/-^* and *Igf2*^*ΔGGCT*^ myotubes in comparison to WT cells (Figure 2B). This was associated with increased myosin heavy chain and myogenin expression in *Zbed6^-/-^* and *Igf2*^*ΔGGCT*^ myotubes (Figure 2C and 2D).

**Figure 2.**
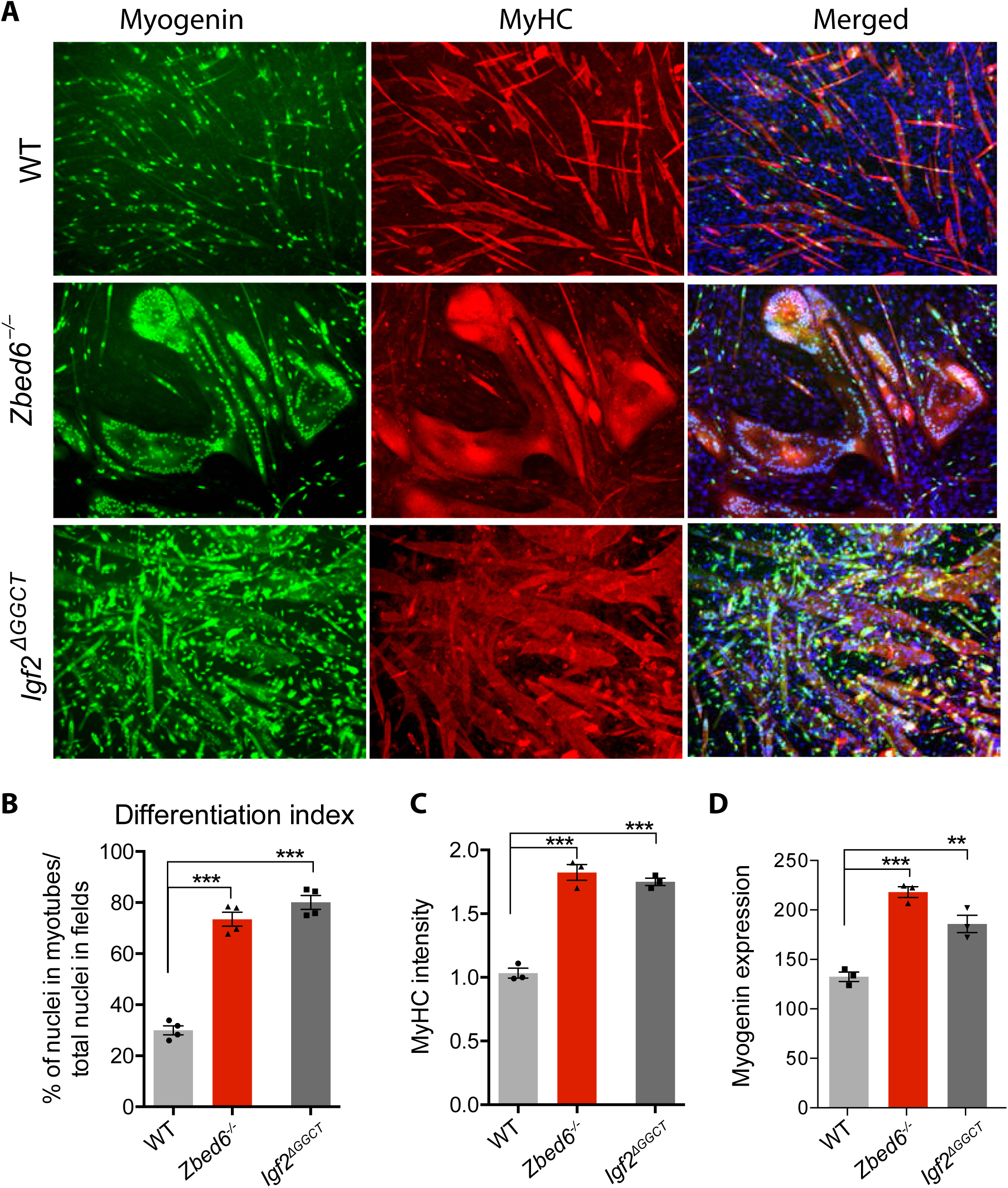
*Zbed6^-/-^* and *Igf2*^*ΔGGCT*^ myoblasts develops hypertrophic myotubes. (A) Immunofluorescence staining of differentiated WT, *Zbed6^-/-^* and *Igf2*^*ΔGGCT*^ myotubes using anti-myogenin antibody (green), anti-myosin-heavy chain (MyHC) antibody (red) and DAPI (blue). (B) Differentiation index of WT, *Zbed6^-/-^* and *Igf2*^*ΔGGCT*^ myotubes, calculated as the percentage of nuclei in myotubes to the total number of nuclei in the same field (Filigheddu et al., 2007). (C) The relative intensity of MyHC staining in the differentiated myotubes. (D) qPCR analysis of *Myogenin* expression in myotubes, the graph present the relative expression to WT myoblast (mean±SEM), **P<0.01, ***P<0.001, Student’s t-test.

### Quantitative SILAC proteomic and transcriptomic analyses of *Zbed6^-/-^* and *Igf2*^Δ^*GGCT* myoblasts

To determine the possible transcriptional targets of ZBED6 during myogenesis, we performed both transcriptomic and mass spectrometry (MS)-based quantitative proteomic analyses of *Zbed6^-/-^*, *Igf2*^*ΔGGCT*^ and WT myoblasts. Differentially regulated genes/proteins and pathways were analyzed with a special focus on genes that showed differential expression in both *Zbed6^-/-^* and *Igf2*^*ΔGGCT*^ myoblasts in order to explore to which extent the observed changes in gene expression in *Zbed6^-/-^* clones are secondary effects due to increased IGF2 expression.

The Stable Isotope Labeling with Amino Acids in Cell Culture (SILAC)-MS technique was used to quantify the changes in the total proteome of the mutant myoblasts. Quantification analysis using MaxQuant identified around 4,000 proteins in each cell line with at least two unique peptides detected in each replicate. The differential expression (DE) analysis of SILAC data showed 312 and 855 DE proteins in *Zbed6^-/-^* medium and lysate fractions, respectively, and 220 and 350 DE proteins in *Igf2*^*ΔGGCT*^ medium and lysate fractions, respectively (*P*<0.05, after Benjamini–Hochberg correction for multiple testing), (Figure S1, Table S1). The transcriptome analysis of *Zbed6^-/-^*, *Igf2*^*ΔGGCT*^ and WT myoblasts identified around 12,000 expressed genes with at least one read count per million (cpm). DE analysis of transcriptome data revealed ~3,000 DE genes in *Zbed6^-/-^* and ~2,500 in *Igf2*^*ΔGGCT*^ myoblasts (Table S2). We integrated the SILAC and RNA-seq data in order to explore the correlation between changes in mRNA and protein expression, to gain further understanding of how the ZBED6-*Igf2* axis affects myoblasts. We detected 381 and 196 genes to be DE in both SILAC and RNA-seq in *Zbed6^-/-^* and *Igf2*^*ΔGGCT*^ myoblasts, respectively. Moreover, we found a significant positive correlation between DE genes and proteins in *Zbed6^-/-^* (r=0.49) and *Igf2*^*ΔGGCT*^ (r=0.55) myoblasts (Figure 3A). Strikingly, the dramatic up-regulation of *Igf2* was detected in both *Zbed6^-/-^* and *Igf2*^*ΔGGCT*^ myoblasts at both the transcriptome and proteome level (Figure 3A).

**Figure 3.**
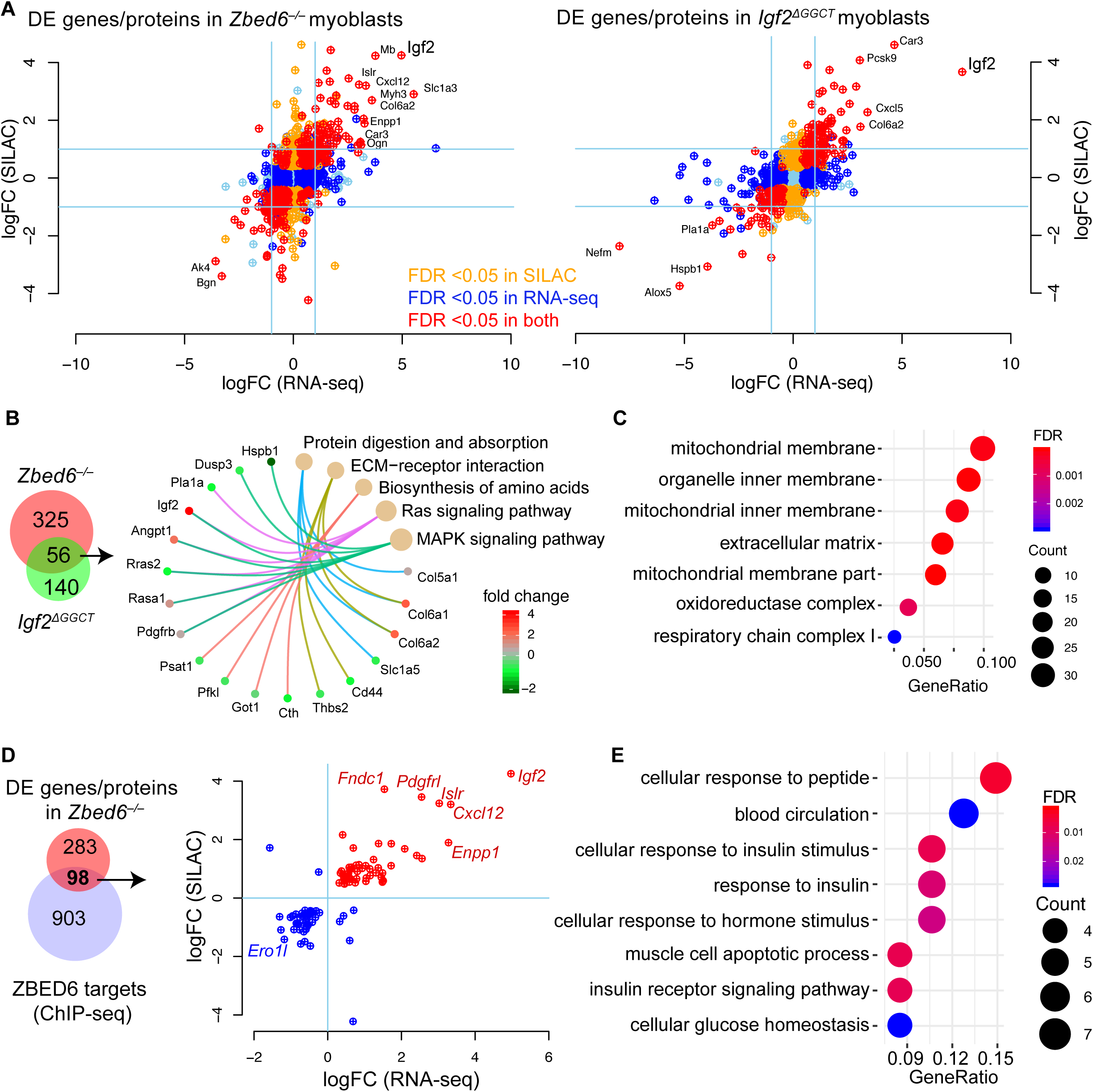
SILAC proteomic and transcriptome analyses of *Zbed6^-/-^* and *Igf2*^*ΔGGCT*^ myoblasts. (A) Expression of identified proteins and genes by SILAC and RNA-seq data in *Zbed6^-/-^* (left) and *Igf2*^*ΔGGCT*^ (right) myoblasts. The values are presented as log fold change (logFC) to WT cells and colored based on the FDR<0.05 values. (B) Intersection of DE proteins in both *Zbed6^-/-^* and *Igf2*^*ΔGGCT*^ myoblasts (left), KEGG pathway analysis of the shared 56 DE proteins (right). (C) GO analysis of the 325 DE proteins in *Zbed6^-/-^* myoblasts. GeneRatio indicates the number of genes found in each term as a proportion of the total number of examined genes. (D) Intersection of DE proteins in *Zbed6^-/-^* myoblasts and putative ZBED6 targets that are expressed in C2C12 cells and detected by SILAC and RNA-seq (left), the fold changes of genes/proteins with ZBED6 binding sites (left). (E) GO analysis of up-regulated genes/proteins in *Zbed6^-/-^* myoblasts with ZBED6 binding sites.

In order to distinguish between the DE genes caused by ZBED6 inactivation and those that are secondary due to *Igf2* up-regulation, we calculated the overlap between DE genes and proteins in both *Zbed6^-/-^* and *Igf2*^*ΔGGCT*^ myoblasts. This analysis showed 56 shared DE proteins in *Zbed6^-/-^* and *Igf2*^*ΔGGCT*^ cells, and 325 DE proteins unique to *Zbed6^-/-^* myoblasts, while 140 was unique to *Igf2*^*ΔGGCT*^ cells (Figure 3B, left). KEGG pathway analysis of those 56 shared DE proteins revealed a significant enrichment of proteins involved in extra cellular matrix (ECM)−receptor interaction, and the MAPK and RAS signaling pathways (Figure 3B, right). Interestingly, the cellular component analysis of DE proteins unique to *Zbed6^-/-^* myoblasts showed a significant enrichment for mitochondrial membrane proteins (Figure 3C), while a similar analysis for the 56 shared DE proteins and the DE proteins unique to *Igf2*^*ΔGGCT*^ myoblasts showed an enrichment for extracellular matrix proteins, and not for mitochondrial terms (Figure S2).

Our previous ChIP-seq analysis for ZBED6 identified thousands of putative target genes in C2C12 myoblasts (Markljung et al., 2009; Jiang et al., 2014). Here, we combined the SILAC, RNA-seq and ChIP-seq data to find out functional targets for ZBED6. The previously described ChIP-seq peaks in C2C12 cells were associated with about 3,000 genes, i.e. about 15% of the genes in the mouse genome. As many as 1,001 of the about 4,000 proteins (25%) detected in the SILAC analysis corresponded to a gene associated with a ZBED6 ChIP-seq peak. This highly significant overrepresentation (P<0.001) is consistent with the notion that ZBED6 binds open chromatin (Jiang et al., 2014). Since 25% of the proteins detected by SILAC corresponded to a gene associated with a ChIP-seq peak it is expected that 25% (95) of the 381 genes detected as differentially expressed both at the mRNA and protein level due to chance only. We found 98 genes in our data (Figure 3D, left). This implies that we cannot draw any firm conclusion on true ZBED6 targets in mouse C2C12 cells, other than the well-established *Igf2* locus, based on these data. In fact, *Igf2* is the gene showing the most striking differentially expressed gene between WT and *Zbed6^-/-^* cells (Figure 3D, right). However, five other genes, highlighted in Figure 3D, showed a striking up-regulated expression after silencing of the ZBED6 repressor suggesting that they may be functional targets. The gene ontology (GO) analysis of the transcripts with significant up-regulation in *Zbed6^-/-^* myoblasts and associated with ChIP-seq peaks showed a significant enrichment for proteins involved in cellular response to insulin stimulus (Figure 3E). These genes include *Insulin Like Growth Factor 1 Receptor* (*Igf1r*), *Phosphoinositide-3-Kinase Regulatory Subunit 1 (Pik3r1)*, and *Ectonucleotide pyrophosphatase phosphodiesterase 1* (*ENPP1*), a negative modulator of insulin receptor (IR) activation.

### ZBED6 modulates transcriptional regulation of differentiated myotubes partially through IGF2 signaling

To investigate the possible role of the ZBED6-*Igf2* axis on transcriptional regulation during myogenesis, we performed transcriptome analyses of *Zbed6^-/-^*, *Igf2*^*ΔGGCT*^ and WT cells after differentiation into myotubes. First, we analyzed the differentially expressed (DE) genes in WT myoblasts vs. myotubes to explore the overall transcriptional changes during myotube formation. The counting of aligned reads using the STAR tool (Dobin et al., 2013) identified ~12,000 expressed genes in myoblasts with at least one read count per million (cpm) (Figure S3A). The expression of ~3,900 genes was found to be changed significantly, with 2,200 up-regulated and 1,700 down-regulated after differentiation into myotubes (*P*<0.05, after Benjamini–Hochberg correction for multiple testing), (Figure S3B). GO analysis of up-regulated genes revealed a significant enrichment of cell adhesion, muscle proteins and muscle contraction genes, while the down-regulated genes were enriched for cell cycle and mitotic nuclear division categories (Figure S3C).

Transcriptome analysis comparing WT and *Zbed6^-/-^* cells after differentiation identified 2,673 DE genes (log fold change >0.5; *P*<0.05, after Benjamini–Hochberg correction for multiple testing), with 1,243 up-regulated and 1,430 down-regulated genes. Furthermore, 2,630 genes showed a significant differential expression in the comparison of *Igf2*^*ΔGGCT*^ and WT cells after differentiation, with 1,278 up-regulated and 1,352 down-regulated in mutant cells (Table S4). The dissection of the DE genes based on the direction of the change revealed a highly significant 30% overlap between DE genes in *Zbed6^-/-^* and *Igf2*^*ΔGGCT*^ myotubes (Figure 4A).

**Figure 4.**
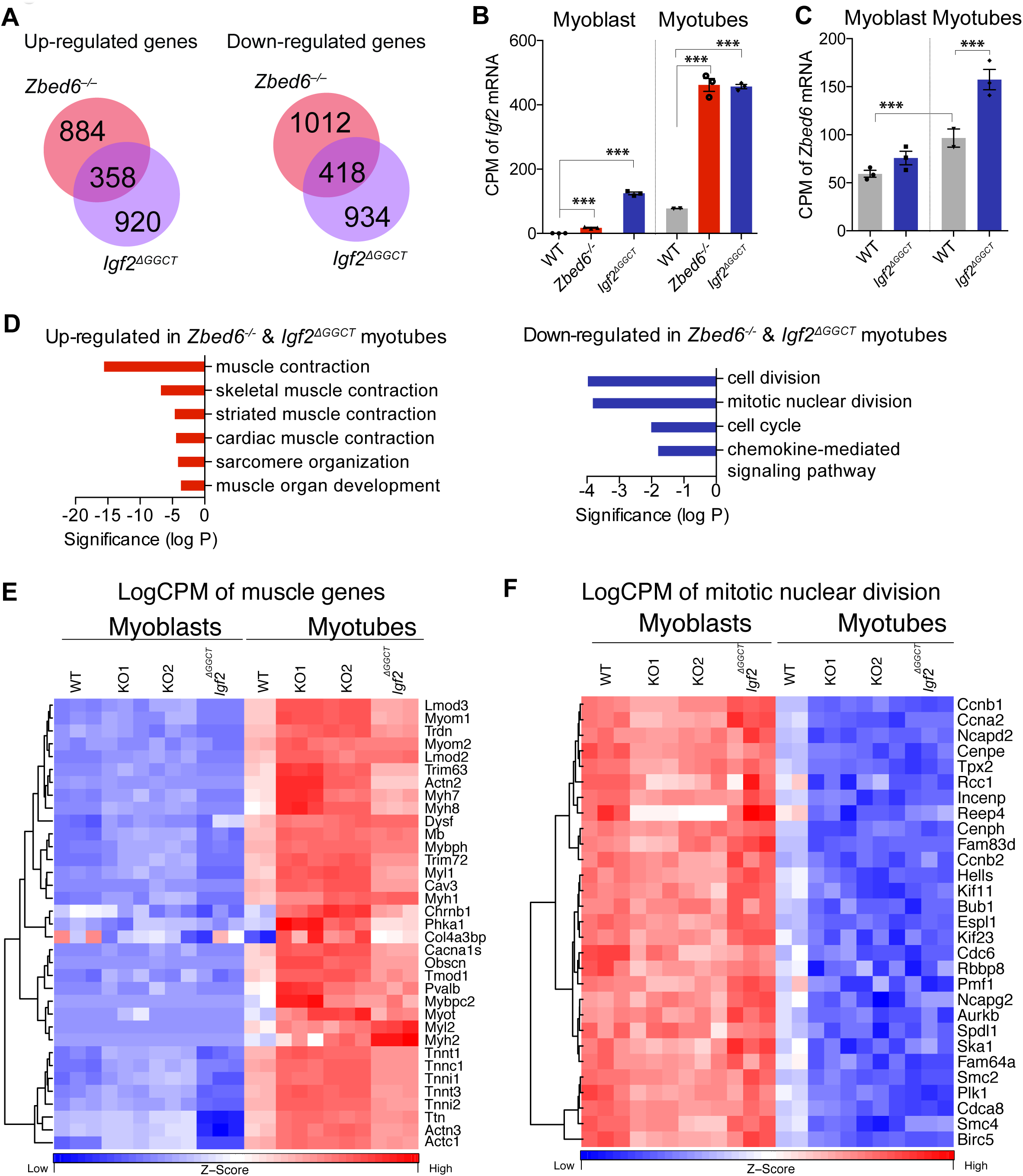
Transcriptome analysis of *Zbed6^-/-^* and *Igf2*^*ΔGGCT*^ myotubes. (A) Intersection of up-regulated and down-regulated DE genes in *Zbed6^-/-^* myotubes (red) vs. *Igf2*^*ΔGGCT*^ myotubes (blue). (B and C) Expression analysis of *Igf2* (B) and *Zbed6* (C) mRNA in WT, *Zbed6^-/-^* and *Igf2*^*ΔGGCT*^ myoblasts and myotubes as count per million (CPM). (D) GO analysis of up-regulated (left) and down-regulated (right) DE genes in both *Zbed6^-/-^* and *Igf2*^*ΔGGCT*^ myotubes. Bars show multiple testing corrected *P*-value for enriched GO categories. (E) Heatmap of muscle-specific genes that were found in muscle contraction GO categories. Expression values are presented as logCPM and color-scaled from blue (low-expression) to red (high-expression). Each column represents an individual sample of *Zbed6^-/-^* (KO1 and KO2), *Igf2*^*ΔGGCT*^ and WT groups. (F) Heatmap of genes that were found in cell cycle and mitotic nuclear division GO categories.

The expression of *Igf2* is known to be upregulated during myoblast differentiation (Florini et al., 1991). In this study, we detected the same pattern as *Igf2* was up-regulated 100-fold after differentiation of WT cells (Figure 4B). Furthermore, the *Igf2* mRNA expression was up-regulated 500-fold in *Zbed6^-/-^* and *Igf2*^*ΔGGCT*^ myotubes in comparison to WT myoblast and 6-fold in comparison to WT myotubes (Figure 4B). Interestingly, this up-regulation of *Igf2* expression was accompanied with a significant but modest increase in the expression of the endogenous *Zbed6* mRNA (Figure 4C), which indicates a positive correlation between *Igf2* and *Zbed6* expression.

The GO analysis of the up-regulated genes in both *Zbed6^-/-^* and *Igf2*^*ΔGGCT*^ myotubes exhibited a striking enrichment of muscle-specific genes (Figure 4D, left). This included genes for myosin heavy and light chains (*Myl1, Myl2, Myh1, Myh2, Myh7* and *Myh8*), troponins (*Tnnt1, Tnnt3, Tnnc1* and *Tnni2*), myomesins (*Myom1* and *Myom2*), alpha actinin (*Actn2 and Actn3*), leiomodin (*Lmod2* and *Lmod3*), titin (*Ttn*) and myoglobin (*Mb*). In contrast, genes involved in cell division and cell cycle regulation such as cyclins (*Ccnb1*, *Ccna2* and *Ccnb2*), were enriched among down-regulated genes in both *Zbed6^-/-^* and *Igf2*^*ΔGGCT*^ myotubes (Figure 4D, right). The expression of genes belonging to these GO categories are presented as logCPM values in myoblast and myotubes (Figure 4E and 4F). The results demonstrate a remarkable shift in their expression before and after differentiation, and how these differences are enhanced in *Zbed6^-/-^* and *Igf2*^*ΔGGCT*^ myotubes.

### ZBED6 over-expression impairs myoblast differentiation

The role of ZBED6 in myogenesis was further investigated by overexpressing ZBED6 (ZBED6-OE) in C2C12 myoblasts and then induce differentiation. Immunofluorescence staining against myogenin and MyHC revealed poor differentiation of ZBED6-OE myoblasts, with a marked reduction in cells expressing myogenin and myosin (Figure 5A). This observation was in stark contrast to the myotube hypertrophy observed in *Zbed6^-/-^* C2C12 cells after differentiation. We performed RNA-seq analysis of ZBED6-OE and control cells after 72h of differentiation. The bioinformatic analysis of differentiated myoblasts revealed 1,560 down-regulated genes and 1,157 up-regulated genes (log fold change >0.5; *P*<0.05, after Benjamini–Hochberg correction for multiple testing) in ZBED6-OE vs. control cells (Figure S4). The most affected genes in response to ZBED6-OE included down-regulation of *Igf2,* myogenin (*Myog*) and myosin heavy chain 3 (*Myh3*) (Figure 5B, figure S4). The GO analysis of down-regulated genes showed a significant enrichment of muscle-specific genes, while the up-regulated genes were primarily related to cell cycle regulation and cell division (Table 1), thus the opposite trend compared with *Zbed6^-/-^* cells. As many as 463 genes were significantly down-regulated in differentiated ZBED6-OE myoblasts and significantly up-regulated in *Zbed6^-/-^* myotubes (Figure 5C). These represents about 40% of the up-regulated genes in *Zbed6^-/-^* myotubes. We examined these genes and the corresponding pathways in more detail. The GO analysis revealed a striking enrichment in muscle-specific categories (Table S5). Among the enriched KEGG pathways, we found cardiac muscle contraction, hypertrophic cardiomyopathy (HCM), and calcium, insulin and AMPK signaling (Table S5). We examined the genes present in the AMPK and insulin signaling pathways and compared their expression in *Zbed6^-/-^, Igf2*^*ΔGGCT*^ and *ZBED6-OE* cells after differentiation. Interestingly, the key components of these pathways were found to be up-regulated in *Igf2*^*ΔGGCT*^ myotubes as well (Figure 5D). For instance, the expression of phosphatidylinositol-4,5-bisphosphate 3-kinase catalytic subunit beta (*Pik3cb*), glycogen synthase 1 (*Gys1*) and AMP-activated protein kinase alpha2 (*Prkaa2*), beta2 (*Prkab2*) and gamma3 (*Prkag3*) subunits were found to be up-regulated in *Zbed6^-/-^* and *Igf2*^*ΔGGCT*^ myotubes, and down-regulated in ZBED6-OE differentiated cells (Figure 5D). Activation of the PI3K pathway and its downstream targets plays a central role in myogenesis. Interestingly, *Prkaa2*, *Prkab2* and *Prkag3*, all up-regulated in *Zbed6^-/-^* and *Igf2*^*ΔGGCT*^ myotubes, encode the α2, β2 and γ3 subunits. These subunits form a specific isoform of AMPK that shows tissue-specific expression in white skeletal muscle (Barnes et al., 2004). The results imply that the interaction between ZBED6-*Igf2* has an essential role in muscle development and influences muscle metabolism.

**Table 1.** GO analysis of up-regulated and down-regulated DE genes in ZBED6-OE vs. control (GFP) differentiated myoblasts

| Term | Count | FDR |
| --- | --- | --- |
| Down-regulated |  |  |
| muscle contraction | 28 | 2.5E-11 |
| cell adhesion | 75 | 1.2E-09 |
| multicellular organism development | 129 | 6.6E-09 |
| cardiac muscle contraction | 24 | 3.1E-08 |
| skeletal muscle contraction | 19 | 4.3E-08 |
| Up-regulated |  |  |
| cell cycle | 127 | 4.7E-25 |
| mitotic nuclear division | 81 | 9.3E-24 |
| cell division | 92 | 1.4E-22 |
| DNA replication | 39 | 7.4E-10 |
| chromosome segregation | 30 | 1.1E-08 |

**Table 2.** GO analysis of up-regulated and down-regulated DE genes in ZBED6-OE vs. control (GFP) proliferating myoblasts

| Term | Count | FDR |
| --- | --- | --- |
| Down-regulated |  |  |
| mRNA processing | 63 | 3.7E-04 |
| DNA replication initiation | 13 | 6.3E-04 |
| **cell cycle** | **98** | **1.2E-03** |
| DNA replication | 34 | 1.6E-03 |
| mitotic nuclear division | 54 | 8.0E-03 |
| Up-regulated |  |  |
| cellular response to interferon-beta | 21 | 1.5E-09 |
| immune system process | 61 | 5.3E-08 |
| innate immune response | 54 | 7.7E-07 |
| defense response to virus | 34 | 1.7E-06 |
| oxidation-reduction process | 86 | 4.8E-03 |

**Figure 5.**
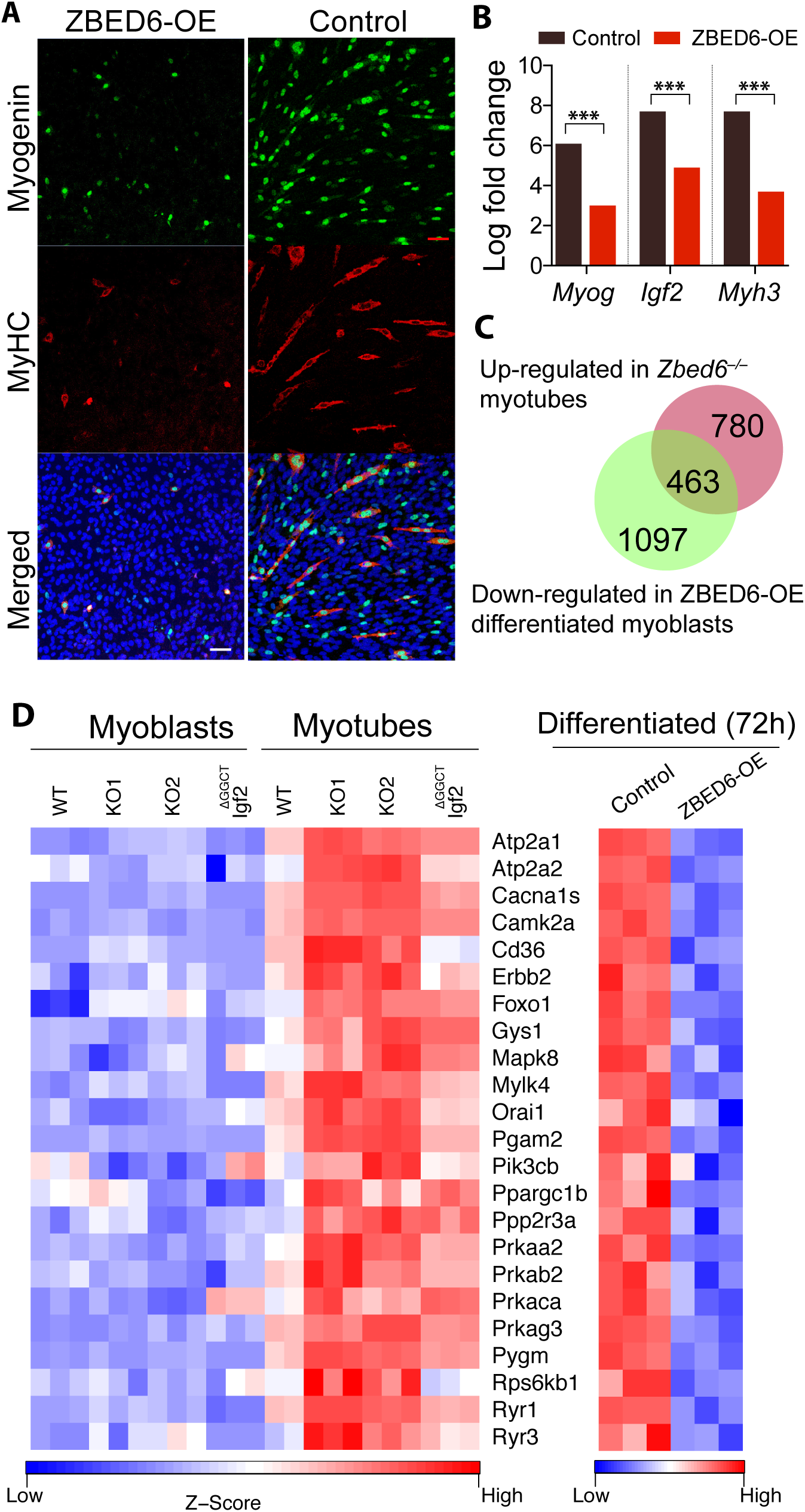
Over-expression of ZBED6 impairs myotube differentiation. (A) Immunofluorescence staining of myogenin and MyHC in 72 h differentiated myoblasts transiently over-expressing ZBED6 (ZBED6-OE). (Scale bar: 50 μm). (B) Log-fold change in the expression of *Myog*, *Igf2* and *Myh3* mRNA in control and ZBED6-OE differentiated myoblasts in comparison to un-differentiated control myoblasts (\*\*\**FDR*<0.001). (C) Intersection of up-regulated DE genes in *Zbed6^-/-^* myotubes (red) vs. down-regulated DE genes in ZBED6-OE differentiated myoblasts (green). (D) Heatmap of genes found in AMPK and insulin signaling pathway.

### Over-expression of ZBED6 in non-differentiated C2C12 cells results in reduced cell viability and cell cycle arrest

Since deletion of ZBED6 promotes cell proliferation and myogenesis, we investigated whether over-expression of ZBED6 causes the opposite effect. Unfortunately, we did not succeed in our attempts to establish a stable myoblast cell line overexpressing ZBED6 suggesting that overexpression may be lethal in C2C12 cells, which is consistent with the results of a previous study (Butter et al., 2010). Therefore, we measured cell viability after transient overexpression of ZBED6 (ZBED6-OE) in C2C12 cells. We found a significant reduction in cell viability in ZBED6-OE cells in comparison to control cells (Figure S5). The cell apoptosis analysis using flow cytometry revealed a significant reduction in the number of live cells in ZBED6-OE (Figure 6A). Moreover, cell cycle analysis using flow cytometry displayed a significant difference in the proportion of cells in different cell cycle phases, in which 82% of ZBED6-OE cells appeared to be in G0/G1 phase and 10% in S-phase, whereas the corresponding proportions in control cells were 58% and 35%, respectively (Figure 6B). These phenotypic changes in ZBED6-OE myoblasts were in complete agreement with the RNA-seq analysis that revealed a significant enrichment of genes involved in cell cycle and cell division processes among the down-regulated genes after ZBED6-OE in proliferating myoblasts (Table 2). Out of 98 down-regulated cell cycle–related genes, 21 were previously identified as putative direct targets of ZBED6 (Jiang et al., 2014; Markljung et al., 2009) (Figure 6C). Some of the putative direct targets with essential role in cell cycle regulation were validated by qPCR (Figure S6). These included the genes for E2f1 and E2f2, members of the E2f family that has an essential role in regulating cell proliferation and controls the transition from G1 to S phase (Wu et al., 2001). There was a striking upregulation of genes involved in immune defense after ZBED6-OE in proliferating myoblasts (Table 2). These results suggest that ZBED6 inhibits proliferation and promotes immune defense in C2C12 cells.

**Figure 6.**
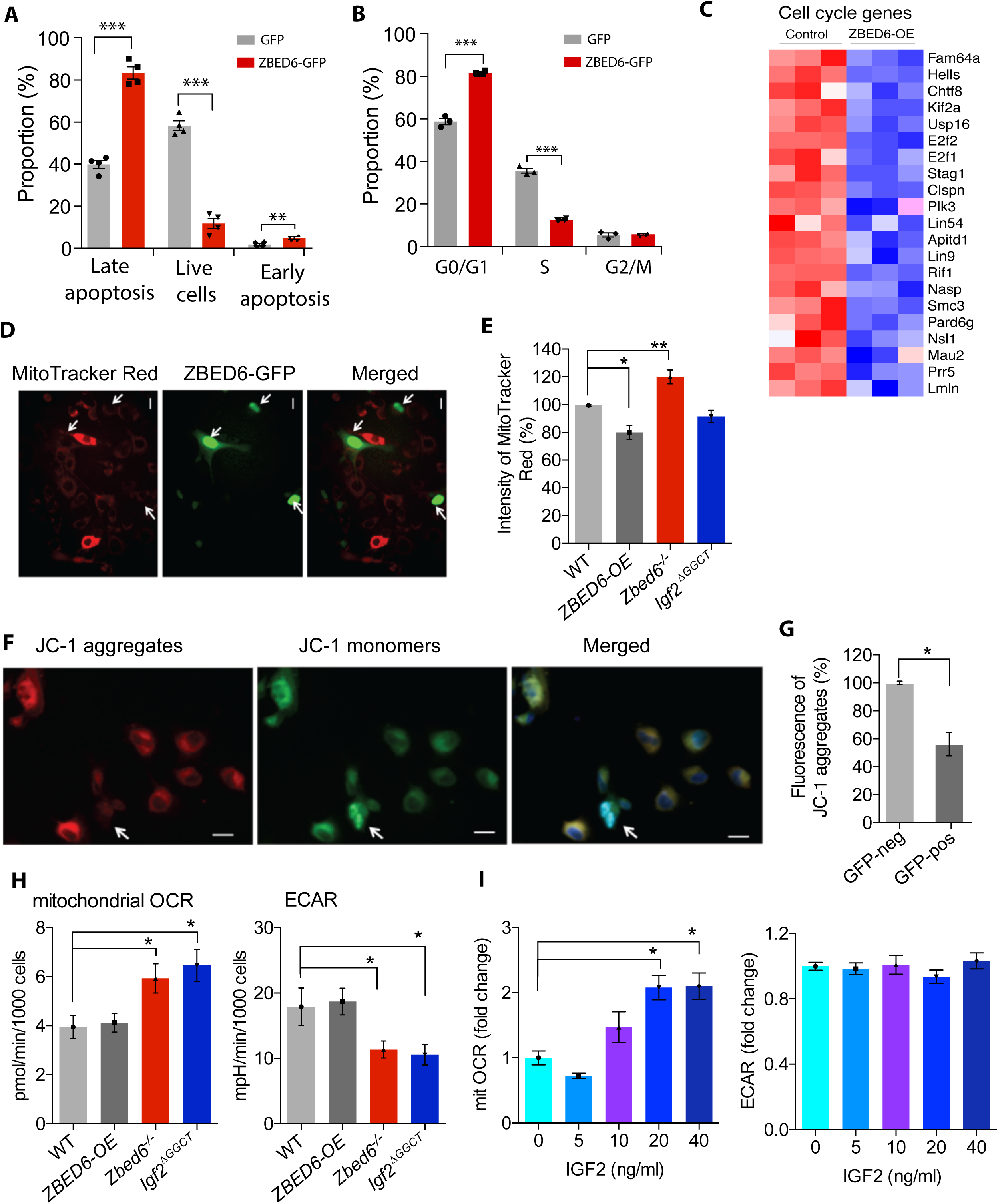
Over-expression of ZBED6 leads to reduced cell viability, cell cycle arrest and reduced mitochondria activity. (A) Cell apoptosis assay of myoblasts over-expressing ZBED6-GFP fusion protein vs. the control myoblasts expressing GFP. (B) Cell cycle analysis of myoblasts over-expressing ZBED6-GFP fusion protein vs. GFP expressing cells. (C) The expression of down-regulated genes found in cell cycle GO categories and containing the consensus ZBED6 binding motif within 1kb of their TSS (Jiang et al., 2014; Markljung et al., 2009). (D) MitoTracker Red labeling of ZBED6 transient overexpressing (ZBED6-GFP) cells. (E) Flow cytometry analysis of the intensity of MitoTracker Red labeling of active mitochondria in WT, ZBED6-over expression (ZBED6-OE), *Zbed6^-/-^* and *Igf2*^*ΔGGCT*^ cells. (F) A representative image of JC-1 aggregates, JC-1 monomer of ZBED6-GFP cells to measure the mitochondrial hyperpolarized membrane potentials. (G) Quantification of the fluorescence intensity of JC-1 aggregates (red) in WT (GFP-neg) and transient ZBED6-OE (GFP-pos) using ImageJ. (Totally 104 WT and 50 ZBED6-OE cells were quantified from three independent experiments.). (H) The oxygen consumption rate (OCR) and extracellular acidification rate (ECAR) were determined with the Extracellular Flux Analyzer XFe96 in C2C12 cells (WT, ZBED6-OE, *Zbed6^-/-^* and *Igf2*^*ΔGGCT*^). Results are means ± SEM for five independent observations. (I) OCR and ECAR were determined in C2C12 WT cells supplemented with 5, 10, 20, 40 ng/ml IGF2 in culture medium. Results were normalized to control condition and are means ± SEM for at least six replicates in each condition. * Denotes *P*<0.05 vs. WT using one-way ANOVA.

### Changes in mitochondrial activity in response to altered ZBED6 expression in myoblasts

The strong correlation between mitochondrial biogenesis and aerobic metabolism, on one hand, and mesenchymal stem cell differentiation, on the other, is well established (Antico Arciuch et al., 2012; Duguez et al., 2004; Hsu et al., 2016). The marked increase in mitochondrial activity that occurs during mesenchymal differentiation is driven by the transcription factor PGC-1α, and IGF2 has been suggested to participate in this process (Lee et al., 2015). Moreover, RNA-seq and SILAC proteomic analyses showed a significant enrichment for mitochondrial membrane proteins among DE proteins in *Zbed6^-/-^* myoblasts (Figure 3C). These observations encouraged us to look closer at mitochondrial activities in response to ZBED6-overexpression or ablation. Flow cytometry analysis of MitoTracker Red intensity, a dye that labels active mitochondria in living cells, indicated a significant reduction in mitochondrial mass in ZBED6-OE cells and an increase in mitochondria in *Zbed6^-/-^* cells, while no change was observed in *Igf2*^*ΔGGCT*^ cells (Figure 6D and 6E). As MitoTracker Red labeling only gives a very crude estimate of mitochondrial mass and activity, we also stained transiently ZBED6-GFP transfected C2C12 cells with JC-1, a probe that gives an estimate of the inner mitochondrial membrane potential. These experiments demonstrated that ZBED6-GFP overexpressing cells displayed lower JC-1 aggregates (red fluorescence) to JC-1 monomers (green fluorescence) ratio (Figure 6F and 6G), indicating a decreased mitochondrial membrane potential in response to overexpressed ZBED6. Both the MitoTracker and JC-1 experiments are in agreement with the SILAC data, and suggest an inhibitory role of ZBED6 on mitochondrial mass/function. Since both *Zbed6^-/-^* and *Igf2*^*ΔGGCT*^ myoblasts exhibited similar phenotypic characteristics regarding growth and differentiation, we explored another part of mitochondrial function in these cells. We analyzed C2C12 cell mitochondrial oxidation rates (OCR) using the Seahorse technique (Malmgren et al., 2009). The OCR and extracellular acidification rate (ECAR) assay revealed an increased OCR (Figure 6H, left) and reduced ECAR in *Zbed6^-/-^* and *Igf2*^*ΔGGCT*^ myoblasts (Figure 6H, right) compared to WT cells. OCR and ECAR were unaffected in ZBED6-OE cells (Figure 6H). As both *Zbed6^-/-^* and *Igf2*^*ΔGGCT*^ myoblasts show increased *Igf2* expression, we hypothesized that the increase in respiration might be an IGF2 effect. To test this hypothesis, we treated cells with recombinant IGF2 and measured the OCR and ECAR dose-response in C2C12 WT cells. We found a 2-fold increase in respiration rates at the higher IGF2 concentrations (20 and 40 ng/ml IGF2) (Figure 6I, left), while no changes were detected in the extracellular acidification rates (Figure 6I, right). Thus, it appears that ZBED6 controls myoblast mitochondrial biogenesis/activity partially via IGF2.

## DISCUSSION

This study conclusively demonstrates that the interaction between ZBED6 and its binding site in *Igf2* plays a critical role in regulating the development of myogenic cells. This conclusion is based on (i) MS quantitative proteomics and transcriptomic analyses, (*ii*) the altered growth rate of myoblasts, (*iii*) effects on myotube formation and maturation, and (*iv*) assessment of mitochondrial activities. Interestingly, the disruption of the ZBED6 binding site in *Igf2* was sufficient to obtain very similar phenotypic effects as observed in the *Zbed6* knock out demonstrating that the phenotypic effects caused by *Zbed6* inactivation in myoblast cells are largely mediated through the regulation of *Igf2* expression.

The initial development of skeletal muscle occurs prenatally and involves the proliferation of myoblasts, which then exit the cell cycle and start differentiation to form myotubes (Dunglison et al., 1999; Stockdale, 1992). It has been reported that the number of myoblasts prenatally greatly influences muscle growth postnatally since the number of muscle fibers are fixed at birth (Rehfeldt et al., 1993; Velloso, 2008). This is of particular interest, since ZBED6 inactivation or disruption of the interaction between ZBED6 and the binding site in *Igf2* promotes the proliferation and growth of myoblasts (Figure 1E, F). The *Zbed6^-/-^* and *Igf2*^*ΔGGCT*^myoblasts had a ~30-fold up-regulation in *Igf2* mRNA expression. This massive increase in *Igf2* expression was rescued towards the levels found in WT cells when ZBED6 was reintroduced in *Zbed6^-/-^* myoblasts, while no changes were observed in case of *Igf2*^*ΔGGCT*^ myoblasts. Thus, ZBED6 represses *Igf2* expression through the binding site located in *Igf2* intron 1.

Our results on myoblast differentiation revealed that *Zbed6^-/-^* and *Igf2*^*ΔGGCT*^ myoblasts were prone to develop mature, hypertrophic and contractile myotubes, with a striking increase in the expression of well-known markers of muscle differentiation, *Igf2*, myogenin and myosin heavy chain (MyHC). In contrast, over-expression of ZBED6 blocked the differentiation of myoblasts, and the expression of *Igf2*, myogenin and MyHC were greatly down-regulated. These phenotypic changes are in agreement with our transcriptome data that revealed a significant ~30% overlap between differentially expressed genes in *Zbed6^-/-^* and *Igf2*^*ΔGGCT*^ myotubes (Figure 4A). Gene ontology analysis of the up-regulated genes in common between *Zbed6^-/-^* and *Igf2*^*ΔGGCT*^ myotubes demonstrated a significant enrichment of muscle-specific categories, including genes encoding myosin heavy and light chains, troponins, titin, myomesins, alpha actinin, leiomodin, and myoglobin. Interestingly, ~40% of the up-regulated DE genes in *Zbed6^-/-^* myotubes were found to be down-regulated in differentiated C2C12 cells after overexpressing ZBED6. The GO analysis of those genes showed an enrichment in muscle-specific categories very similar to what we found among up-regulated genes in *Zbed6^-/-^* myotubes.

SILAC quantitative proteomic analysis of mutant myoblasts revealed a significant enrichment of mitochondrial membrane proteins exclusively among the DE proteins found in *Zbed6^-/-^* but not *Igf2*^*ΔGGCT*^ myoblast (Figure 3C). This observation was confirmed by the decreased mitochondrial membrane potential in response to ZBED6 overexpression (Figure 6D-G). However, the assessment of mitochondrial respiration rate indicated a positive correlation between oxygen consumption rate and the amount of IGF2 protein in myoblasts. This was concluded by the consistent changes in oxygen consumption in *Zbed6^-/-^* and *Igf2*^*ΔGGCT*^ myoblasts, and after the addition of recombinant IGF2 to the growth medium of wild-type myoblasts. Our findings fit well with what has been reported in literature about the essential role of mitochondrial activities in proliferation and differentiation of myoblasts. For instance, respiration-deficient human myoblasts were growing slower than control cells, exhibited low ATP synthesis and demonstrated sever deficiency in myotube formation (Herzberg et al., 1993). Furthermore, it has been indicated that the basal mitochondrial respiration rate was increased one-fold and the maximal respiration increased four-fold in differentiated myotubes (Remels et al., 2010).

We have previously reported that ZBED6 has ~2,500 binding sites all over the genome (Jiang et al., 2014; Markljung et al., 2009). Here we find that the disruption of only one of these binding sites, located in *Igf2* intron 1, resulted in similar phenotypic changes as observed by complete inactivation of *Zbed6* in C2C12 cells. The results suggest that the regulation of IGF2 expression may be the most important role of ZBED6 in skeletal muscle cells, which is consistent with the initial observation that a mutation of this binding site is causing an altered body composition in pigs selected for meat production (Van Laere et al., 2003) and our recent characterization of *Zbed6^-/-^* and *Igf2* knock-in mice (Younis et al., 2018b). However, ZBED6 is essentially found in all cell types and throughout development, so it may interact with other important targets in other cells or during other stages of development.

In summary, we have shown that the up-regulation of *Igf2* expression obtained either by ZBED6 ablation or the deletion of its binding site in intron one of *Igf2*, has an essential role in modulating the metabolism of myogenic cells and promotes differentiation of myoblast cells partially through increasing respiration rate of the mitochondria. This study suggests that the interaction between ZBED6-*Igf2* may be an important therapeutic target for human diseases where anabolism is impaired.

## METHODS

### Cell culture

Mouse myoblast C2C12 cells were obtained from ATCC (CRL-1772), and it is a subclone of the previously established mouse myoblast cell line (Yaffe and Saxel, 1977). The cells were maintained in Dulbecco's Modified Eagle Medium (DMEM) with 2 mM L-glutamine, 1 mM sodium pyruvate and 4.5 g/L glucose (ATCC 30-2001), supplemented with 10% (v/v) heat-inactivated fetal bovine serum (FBS) and penicillin (0.2 U/mL)/streptomycin (0.2 μg/ml)/L-glutamine (0.2 μg/ml) (Gibco) at 37°C in a 5% CO_2_ humidified atmosphere. Differentiation was induced by replacing FBS with 2% horse serum (Gibco). The differentiation medium was changed every 48h. The differentiated myotubes were collected by adding 0.05% Trypsin-EDTA (Gibco) for 1 min at 37°C, which was sufficient to detach the mature myotubes from the plate.

### Genome editing

The coding sequence of *Zbed6* and its binding site in *Igf2* were targeted in C2C12 cells using CRISPR/Cas9 tools. Two specific guide RNAs (gRNA) for *Zbed6* and one for *Igf2* were designed using the CRISPRdirect tools (Naito et al., 2015). The gRNAs sequences were cloned into the Cas9 expressing plasmid pSpCas9(BB)-2A-GFP (PX458), (Addgene plasmid #48138) and co-transfected with linear hygromycin marker (Clontech) into C2C12 cells at passage number 5. Wild-type cells were transfected with empty pSpCas9(BB)-2A-GFP plasmid and linear hygromycin. Transfected cells were kept under selective medium for two weeks. Single-cell clones were screened for a 2.5 kb deletion in *Zbed6* using primers flanking the targeted site (Figure 1A). The *Igf2* targeted clones were screened using primers flanking the targeted site, followed by Sanger sequencing of individual clones.

### Immunofluorescence staining

Cells were cultured in an 8-well slide chamber (BD Falcon) overnight to around 60–70% confluence. The cells were washed with PBS and fixed with 4% (v/v) paraformaldehyde for 10 min at room temperature. The fixed cells were permeabilized with 0.25% (v/v) Triton X-100 and then blocked for non-specific binding with 2% (w/v) BSA in PBS. The primary antibodies for myogenin and myosin heavy chain (Santa Cruz Biotechnology) were diluted 1:500 in PBS containing 1% (w/v) BSA and incubated with the cells overnight. Cells were washed three times with PBS and then incubated with Alexa Flour-labeled secondary antibodies. DAPI was used as counter staining. Slides were analyzed using a confocal microscope (Zeiss LSM 700).

### Real-time quantitative PCR

Total RNA was extracted from cells using the RNeasy Mini kit (Qiagen), including the DNase I treatment. The High Capacity cDNA Reverse Transcription Kit (Applied Biosystems) was used to generate cDNA from the extracted RNA. Quantitative PCR (qPCR) analysis was performed using ABI MicroAmp Optical 384-well Reaction plates on an ABI 7900 real-time PCR instrument (Applied Biosystems). The qPCR was performed using TaqMan Gene Expression Assays that consisted of forward and reverse primers with TaqMan minor groove binder (MGB) probe for each gene (*Zbed6*: Mm04178798_s1, *Igf2*: Mm00439564_m1, *18S*: Mm03928990_g1, Applied Biosystems); 18S and was used as housekeeping gene. For *Myog* and the validated DE genes, the forward and reverse primers (Tables S6) were mixed with SYBR Green Gene Expression Master Mix (Applied Biosystems) in 10 μl total reaction volume.

### Immunoblot analysis

Total protein lysates were prepared using RIPA lysis buffer containing protease inhibitors (Complete Ultra Tablets, Roche). Equal amounts of total lysates were separated by SDS-PAGE (4–15%, Bio-Rad) and transferred to PVDF membranes (Millipore). StartingBlock buffer (Thermo Scientific) was used to block the membrane before the primary anti-ZBED6 antibody (1:1000) was added (Akhtar et al., 2015). Proteins were visualized and detected by the Odyssey system (LI-COR).

### Stable Isotope Labeling with Amino acids in Cell culture (SILAC)

C2C12 cells were cultured in Dulbecco's modified Eagle's medium (DMEM) for SILAC (Thermo Fisher Scientific) supplemented with 10% dialyzed fetal bovine serum (FBS, MWCO 10 kDa; Thermo Fisher Scientific), 100 U/mL penicillin (Thermo Fisher Scientific), 100 μg/mL streptomycin (Thermo Fisher Scientific), 0.25 μg/mL amphotericin B (Thermo Fisher Scientific) and light isotopic labels L-arginine-HCl and L-lysine-2 HCl or heavy isotopic labels ^13^C_6_, _15_N4 L-arginine-HCl (Arg-10) and ^13^C_6_, ^15^N_2_ L-lysine-2 HCl (Lys-8) (Thermo Fisher Scientific). Cells were kept in a humidified atmosphere with 5% CO_2_ at 37°C. To avoid contamination of light amino acids, sub-culturing was performed using Cell dissociation buffer (Thermo Fisher Scientific) instead of trypsin. Isotopic incorporation was checked using a script in R as previously described (Stöhr and Tebbe, 2011) after approximately five cell divisions to confirm complete (>95%) labeling. Arginine-to-proline conversion was assessed by calculating the percentage of heavy proline (Pro-6) containing peptides among all identified peptides, and kept at <5%. Confluent (~80%) cells were washed five times in PBS, and incubated 12 h in serum-free medium. Medium was collected, centrifuged and filtered through a 0.2 μm filter. Protein concentration was measured with a Coomassie (Bradford) assay kit (Thermo Fisher Scientific). Cells were harvested and lysates prepared using M-PER mammalian protein extraction reagent (Thermo Fisher Scientific), and protein concentration was measured using a bicinchoninic acid (BCA) protein assay kit (Thermo Fisher Scientific). Heavy and light cell lysates or media (40 μg each of cell lysate, 170 μg each of medium) were mixed 1:1. Mixed media were subsequently concentrated through spin columns with a cutoff of 3 kDa (Vivaspin, Sartorius). Mixed proteins were separated on a 4-20% Mini-PROTEAN TGX precast gel (Bio-Rad, Hercules, CA). Each gel lane was cut into ten separate pieces, and proteins were reduced in-gel with 10 mM DTT in 25 mM NH_4_HCO_3_, thereafter alkylated with 55 mM iodoacetamide in 25 mM NH_4_HCO_3_, and finally digested with 17 ng/μl sequencing-grade trypsin (Promega) in 25 mM NH_4_HCO_3_ using a slightly modified in-gel digestion protocol (Shevchenko et al., 1996). The produced peptides were eluted from the gel pieces using 1% (v/v) formic acid (FA) in 60% (v/v) acetonitrile, dried down in a vacuum centrifuge (ThermoSavant SPD SpeedVac, Thermo Scientific), and finally dissolved in 1% (v/v) FA. The experiments were run in triplicates, of which one was reciprocal (reverse labeling).

### Liquid chromatography and mass spectrometry

Peptide samples were desalted using Stage Tips (Thermo Fisher Scientific) according to the manufacturer's protocol, and subsequently dissolved in 0.1% (v/v) FA. Samples were separated by RP-HPLC using a Thermo Scientific nLC-1000 with a two-column setup; an Acclaim PepMap 100 (2 cm x 75 μm, 3 μm particles, Thermo Fisher Scientific) pre-column was connected in front of an EASY-Spray PepMap RSLC C18 reversed phase column (50 cm x 75 μm, 2 μm particles, Thermo Fisher Scientific) heated to 35°C, running solvent A (H_2_O and 0.1% (v/v) FA). A gradient of 2–40% solvent B (acetonitrile and 0.1% (v/v) FA) was run at 250 nL/min over a period of 3 h. The eluted peptides were analyzed on a Thermo Scientific Orbitrap Fusion Tribrid mass spectrometer, operated at a Top Speed data-dependent acquisition scan mode, ion-transfer tube temperature of 275°C, and a spray voltage of 2.4 kV. Full scan MS spectra (m/z 400 – 2000) were acquired in profile mode at a resolution of 120,000 at m/z 200, and analyzed in the Orbitrap with an automatic gain control (AGC) target of 2.0e5 and a maximum injection time of 100 ms. Ions with an intensity above 5.0e3 were selected for collision-induced dissociation (CID) fragmentation in the linear ion trap at a collision energy of 30%. The linear ion trap AGC target was set at 1.0e4 with a maximum injection time of 40 ms, and data was collected at centroid mode. Dynamic exclusion was set at 60 s after the first MS1 of the peptide. The system was controlled by Xcalibur software (version 3.0.63.3, Thermo Scientific). Instrument quality control was monitored using the Promega 6×5 LC-MS/MS Peptide Reference Mix (Promega) before and after each MS experiment run, and analyzed using PReMiS software (version 1.0.5.1, Promega).

### Mass spectrometric data analysis

Data analysis of raw files was performed using MaxQuant software (version 1.5.6.5) and the Andromeda search engine (Cox and Mann, 2008; Tyanova et al., 2016), with cysteine carbamidomethylation as a static modification and Arg-10, Lys-8, methionine oxidation and protein N-terminal acetylation as variable modifications. First search peptide MS1 Orbitrap tolerance was set to 20 ppm. Iontrap MS/MS tolerance was set to 0.5 Da. The re-quantify option was enabled to get ratios where only one isotope pattern was found. Match between runs was also enabled, to identify peptides where only MS1 data was available. Minimum label ratio count was set to 2, and the advanced ratio estimation option was enabled. Peak lists were searched against the UniProtKB/Swiss-Prot *Mus musculus* proteome database (UP000000589, version 2016-01-12) with a maximum of two trypsin miscleavages per peptide. The contaminants database of MaxQuant was also utilized. A decoy search was made against the reversed database, where the peptide and protein false discovery rates were both set to 1%. Only proteins identified with at least two peptides of at least 7 amino acids in length were considered reliable. The peptide output from MaxQuant was filtered by removing reverse database hits, potential contaminants and proteins only identified by site (PTMs). Intensity values were first normalized using variance stabilization method, adjusted for batch effect and fitted to linear model (Huber et al., 2002; Ritchie et al., 2015). The empirical Bayes moderated t-statistics and their associated *P*-values were used to calculate the significance of DE proteins (Smyth, 2004). The *P*-values were corrected for multiple testing using the Benjamini–Hochberg procedures (Benjamini and Hochberg, 1995).

### RNA sequencing

Myoblasts or the collected myotubes were washed in PBS and total RNA was extracted using the RNeasy Mini kit (QIAGEN). The RNA quality and integrity was measured with a RNA ScreenTape assay (TapeStation, Agilent Technologies). Strand-specific, 3′ end mRNA sequencing libraries were generated using QuantSeq 3′ mRNA-Seq Library Prep Kit (Lexogen) following the manufacturer's instructions. For each sample, 2 µg of total RNA were poly-A selected using oligo-dT beads to enrich for mRNA, and the RNA-seq libraries were amplified by 12 PCR cycles. The libraries were size-selected for an average insert size of 150 bp and sequenced as 50 bp paired-end reads using Illumina HiSeq. Sequence reads were mapped to the reference mouse genome (mm10) using STAR 2.5.1b with default parameters (Dobin et al., 2013). HTSeq-0.6.1 (Python Package) (Anders et al., 2015) was used to generate read counts and edgeR (Bioconductor package) (Robinson et al., 2009) was used to identify differentially expressed (DE) genes using gene models for mm10 downloaded from UCSC (www.genome.ucsc.edu). The abundance of gene expression was calculated as count-per-million (CPM) reads. Genes with less than one CPM in at least three samples were filtered out. The filtered libraries were normalized using the trimmed mean of M-values (TMM) normalization method (Robinson and Oshlack, 2010). P-values were corrected for multiple testing using the False Discovery Rate (FDR) approach. The DE genes were submitted to The Database for Annotation, Visualization and Integrated Discovery (DAVID, v6.8) (Huang et al., 2008) for gene ontology analysis. All expressed genes in C2C12 cells were used as background, and the Biological Process and KEGG pathway tables were used to identify enriched GO terms.

### Cell growth measurements

WT cells at different passages (P5, P12 and P12), WT cells transfected with the Cas9 plasmid (WT-Cas9), two *Zbed6^-/-^* and two *Igf2*-mutant clones were seeded in 24-well plates (10,000 cells per well) in growth media, and were cultured for 6 days with real-time measurement of cell density every 12 h using an IncuCyte instrument (Essen Bioscience). The experiments were carried out using biological triplicates of each cell line.

### Cell viability assay

Cells were seeded in 24-well plates (30,000 cells/well) and let to attach. A day after, the cells were transfected with GFP (control) or ZBED6-GFP overexpressing constructs (ZBED6-OE). At 24 h post transfection, the cells were incubated with growth media containing 10% (v/v) PrestoBlue (Invitrogen). The reduction of PrestoBlue reagent was measured on a Tecan Sunrise Plate Reader at four time points post incubation (10 min, 30 min, 2 h and 4 h), with the following parameters: bottom-read fluorescence (excitation 560 nm, emission 590 nm).

### Cell cycle

C2C12 cells transiently expressing ZBED6-GFP or GFP were analyzed for cell cycle profile using Click-iT EdU flow cytometry assay kit (Invitrogen) and FxCycleTM violet stain (Invitrogen) for total DNA staining. Cells were incubated with EdU for 2 h according to the manufacturer’s instructions before fixation and DNA staining. Cells were analyzed on LSR Fortessa (BD Biosciences), and results were analyzed using the FACSDiva software (BD Biosciences).

### Cell apoptosis

C2C12 cells were harvested and fixed with 4% PFA (not permeabilized), and controls were fixed and permeabilized with Cytofix/Cytoperm solution (BD Biosciences) before staining with V450-Annexin-V (BDBiosciences) and/or DRAQ7 (Biostatus). Cells were stained with V450-Annexin-V and DRAQ7 according to the manufacturer’s protocol. Cells were incubated for 15-30 min at RT in dark. Samples were analyzed on BD LSRFortessa flow cytometer using BD FACSDiva software. Viable cells were Annexin-V negative and DRAQ7 negative (AnnV-, PI-) staining, while cells in early apoptosis were Annexin-V positive and DRAQ7 negative. Cells in late-apoptosis/necrosis were Annexin-V positive and DRAQ7 positive.

### MitoTracker Red and JC-1 staining

C2C12 cells were stained for active mitochondria by 50nM MitoTracker Red FM (Invitrogen) for 30 min. After washing, the cells were either fixed for confocal imaging by a Zeiss 780 confocal microscope or directly examined for MitoTracker Red intensity using a FACSCalibur flow cytometer (BD Biosciences). Mitochondrial membrane potential of C2C12 cells was semiquantitatively determined by the fluorescent probe JC-1 (4 μM, 30 min incubation, Sigma-Aldrich). After careful washing, the cells were imaged by confocal microscope and the fluorescence of JC-1 aggregates at hyperpolarized membrane potential (585 nm) was quantified by Image J.

### Oxygen consumption and extracellular acidification rates

The oxygen consumption rate (OCR) and extracellular acidification rate (ECAR) were determined using the Extracellular Flux Analyzer XF^e^96 (Seahorse Bioscience). Cells were cultured in 96-well plates (Seahorse Biosciences) in normal culture medium for 24-48 h. After culture, assays were performed in XF assay medium (Seahorse Biosciences) set to pH 7.4 and supplemented with 25 mM glucose. OCR and ECAR were measured during the last 30 min of the 1 h incubation in XF assay medium, which was followed by the injection of inhibitors of electron transport chain, 5 μmol/l rotenone and 5 μmol/l antimycin A, to inhibit the mitochondrial respiration. The remaining OCR was considered as non-mitochondrial respiration. To calculate the mitochondrial respiration, non-mitochondrial OCR was subtracted from the total OCR. Data were normalized to cell number in the wells and presented as pmol/min/10,000 cells. Recombinant mouse IGF2 (R&D Systems, MN, USA), 5-40 ng/ml was supplemented to culture medium for 24 h before measurement.

## ACKNOWLEDGMENTS

**We thank Aris Moustakas for valuable comments on the manuscript.** The work was funded by grants from The Knut and Alice Wallenberg Foundation and Swedish Research Council. Sequencing was performed by the SNP&SEQ Technology Platform, supported by Uppsala University and Hospital, SciLifeLab and Swedish Research Council (80576801 and 70374401). Computer resources were provided by UPPMAX, Uppsala University.

## AUTHOR CONTRIBUTIONS

LA and NW conceived the study. SY was responsible for gene editing and most of the characterization of the different cell lines including RNAseq and bioinformatics analysis. SY, XC and ML conducted the SILAC proteomics of the cell lines. RN carried out qPCR validation, cell cycle and apoptosis analysis. XW, ES, PB and NW were responsible for the characterization of mitochondrial function. SY and LA wrote the paper with input from other authors. All authors approved the manuscript before submission.

## FIGURE LEGENDS

**Figure S1. Volcano plots for differentially expressed proteins in SILAC data of medium and lysate fractions of *Zbed6^-/-^* and *Igf2*^*ΔGGCT*^ myoblasts. The proteins showing the most striking differential expression are highlighted.**

**Figure S2. GO analysis of 56 shared differentially expressed proteins in *Zbed6^-/-^* and *Igf2*^*ΔGGCT*^myoblasts (top) and 140 differentially expressed proteins unique to *Igf2*^*ΔGGCT*^ myoblasts (bottom).**

**Figure S3. Transcriptome analysis of differentially expressed genes in WT differentiated myotubes.** (A) Scatter plot of all expressed genes in WT myoblast vs. myotubes. (B) Volcano plot for differentially expressed genes of WT myoblast vs. myotubes. Red dots indicate significant fold-change (FDR<0.05). The genes showing the most striking differential expression are highlighted, *Zbed6* and *Igf2* are highlighted with blue dots. (C) Gene ontology (GO) analysis of significantly up-regulated (up-reg) DE genes (left) and down-regulated genes (right) in myotubes. Bars show the significance for GO enriched categories.

**Figure S4. Volcano plot for differentially expressed genes (FDR<0.05) in 72 h differentiated myoblasts (Control vs. ZBED6-OE). The genes showing the most striking differential expression are highlighted.**

**Figure S5. Cell viability assay of myoblasts over-expressing ZBED6-GFP fusion protein vs. the control myoblasts expressing GFP.**

**Figure S6. qPCR validation of cell cycle genes associated with ZBED6 binding sites.** (mean±SEM), *P<0.05, **P<0.01, Student’s t-test.

**Table S1. SILAC data for differentially expressed proteins in *Zbed6^-/-^* or *Igf2* ^*ΔGGCT*^ (GGCT) myoblasts**

**Table S2. RNA-seq data for differentially expressed genes in *Zbed6^-/-^* or *Igf2*^*ΔGGCT*^ mutant (GGCT) myoblasts.**

**Table S3. SILAC and RNA-seq data for differentially expressed proteins in *Zbed6^-/-^* myoblasts with ZBED6 binding site identified by ChIP-seq analysis.**

**Table S4. Differentially expressed genes in *Zbed6* knock-out^*-/-*^or *Igf2*^*ΔGGCT*^ differentiated myotubes.**

**Table S5. GO analysis of the differentially expressed (DE) genes after ZBED6-overexpression (ZBED6-OE) in growing (GM) and differentiated (diff) myoblasts.**

**Table S6. The primer sequences used for qPCR analysis.**

## REFERENCES

Akhtar, M., Younis, S., Wallerman, O., Gupta, R., Andersson, L., and Sjöblom, T. (2015). Transcriptional modulator ZBED6 affects cell cycle and growth of human colorectal cancer cells. Proc. Natl. Acad. Sci. 112, 7743–7748.

Anders, S., Pyl, P.T., and Huber, W. (2015). HTSeq-A Python framework to work with high-throughput sequencing data. Bioinformatics 31, 166–169.

Antico Arciuch, V.G., Elguero, M.E., Poderoso, J.J., and Carreras, M.C. (2012). Mitochondrial Regulation of Cell Cycle and Proliferation. Antioxid. Redox Signal. 16, 1150–1180.

Barnes, B.R., Marklund, S., Steiler, T.L., Walter, M., Hjälm, G., Amarger, V., Mahlapuu, M., Leng, Y., Johansson, C., Galuska, D., et al. (2004). The 5′-AMP-activated protein kinase γ3 isoform has a key role in carbohydrate and lipid metabolism in glycolytic skeletal muscle. J. Biol. Chem. 279, 38441–38447.

Benjamini, Y., and Hochberg, Y. (1995). Controlling the False Discovery Rate: A Practical and Powerful Approach to Multiple Testing. J. R. Stat. Soc. Ser. B 57, 289–300.

Butter, F., Kappei, D., Buchholz, F., Vermeulen, M., and Mann, M. (2010). A domesticated transposon mediates the effects of a single-nucleotide polymorphism responsible for enhanced muscle growth. EMBO Rep 11, 305–311.

Cox, J., and Mann, M. (2008). MaxQuant enables high peptide identification rates, individualized p.p.b.-range mass accuracies and proteome-wide protein quantification. Nat. Biotechnol. 26, 1367–1372.

Dobin, A., Davis, C.A., Schlesinger, F., Drenkow, J., Zaleski, C., Jha, S., Batut, P., Chaisson, M., and Gingeras, T.R. (2013). STAR: Ultrafast universal RNA-seq aligner. Bioinformatics 29, 15–21.

Duguez, S., Sabido, O., and Freyssenet, D. (2004). Mitochondrial-dependent regulation of myoblast proliferation. Exp. Cell Res. 299, 27–35.

Dunglison, G.F., Scotting, P.J., and Wigmore, P.M. (1999). Rat embryonic myoblasts are restricted to forming primary fibres while later myogenic populations are pluripotent. Mech. Dev. 87, 11–19.

Filigheddu, N., Gnocchi, V.F., Coscia, M., Cappelli, M., Porporato, P.E., Taulli, R., Traini, S., Baldanzi, G., Chianale, F., Cutrupi, S., et al. (2007). Ghrelin and des-acyl ghrelin promote differentiation and fusion of C2C12 skeletal muscle cells. Mol. Biol. Cell 18, 986–994.

Florini, J.R., Magri, K.A., Ewton, D.Z., James, P.L., Grindstaff, K., Rotwein, P.S., Florinisg, J.R., Magris, K.A., Ewtons, D.Z., Jamesni, P.L., et al. (1991). “Spontaneous” differentiation of skeletal myoblasts is dependent upon autocrine secretion of insulin-like growth factor-II. J. Biol. Chem. 266, 15917–15923.

Herzberg, N.H., Zwart, R., Wolterman, R.A., Ruiter, J.P.N., Wanders, R.J.A., Bolhuis, P.A., and van den Bogert, C. (1993). Differentiation and proliferation of respiration-deficient human myoblasts. Biochim. Biophys. Acta - Mol. Basis Dis. 1181, 63–67.

Hsu, Y.-C., Wu, Y.-T., Yu, T.-H., and Wei, Y.-H. (2016). Mitochondria in mesenchymal stem cell biology and cell therapy: From cellular differentiation to mitochondrial transfer. Semin. Cell Dev. Biol. 52, 119–131.

Huang, D.W., Sherman, B.T., and Lempicki, R.A. (2008). Systematic and integrative analysis of large gene lists using DAVID bioinformatics resources. Nat. Protoc. 4, 44–57.

Huber, W., Von Heydebreck, A., Sültmann, H., Poustka, A., and Vingron, M. (2002). Variance stabilization applied to microarray data calibration and to the quantification of differential expression. In Bioinformatics, (Oxford University Press), pp. S96–S104.

Jiang, L., Wallerman, O., Younis, S., Rubin, C.-J.C.J., Gilbert, E.R.E.R., Sundström, E., Ghazal, A., Zhang, X., Wang, L.L., Mikkelsen, T.S.T.S., et al. (2014). ZBED6 modulates the transcription of myogenic genes in mouse myoblast cells. PLoS One 9, e94187.

Jinek, M., Chylinski, K., Fonfara, I., Hauer, M., Doudna, J.A., and Charpentier, E. (2012). A programmable dual-RNA-guided DNA endonuclease in adaptive bacterial immunity. Science (80-.). 337, 816–821.

Van Laere, A.-S., Nguyen, M., Braunschweig, M., Nezer, C., Collette, C., Moreau, L., Archibald, A.L., Haley, C.S., Buys, N., Tally, M., et al. (2003). A regulatory mutation in IGF2 causes a major QTL effect on muscle growth in the pig. Nature 425, 832–836.

Lee, K.Y., Singh, M.K., Ussar, S., Wetzel, P., Hirshman, M.F., Goodyear, L.J., Kispert, A., and Kahn, C.R. (2015). Tbx15 controls skeletal muscle fibre-type determination and muscle metabolism. Nat. Commun. 6, 8054.

Malmgren, S., Nicholls, D.G., Taneera, J., Bacos, K., Koeck, T., Tamaddon, A., Wibom, R., Groop, L., Ling, C., Mulder, H., et al. (2009). Tight coupling between glucose and mitochondrial metabolism in clonal β-cells is required for robust insulin secretion. J. Biol. Chem. 284, 32395–32404.

Markljung, E., Jiang, L., Jaffe, J.D., Mikkelsen, T.S., Wallerman, O., Larhammar, M., Zhang, X., Wang, L., Saenz-Vash, V., Gnirke, A., et al. (2009). ZBED6, a novel transcription factor derived from a domesticated DNA transposon regulates IGF2 expression and muscle growth. PLoS Biol 7, e1000256.

Naito, Y., Hino, K., Bono, H., and Ui-Tei, K. (2015). CRISPRdirect: Software for designing CRISPR/Cas guide RNA with reduced off-target sites. Bioinformatics 31, 1120–1123.

Ran, F.A., Hsu, P.D., Wright, J., Agarwala, V., Scott, D.A., and Zhang, F. (2013). Genome engineering using the CRISPR-Cas9 system. Nat. Protoc. 8, 2281–2308.

Rehfeldt, C., Fiedler, I., Weikard, R., Kanitz, E., and Ender, K. (1993). It is possible to increase skeletal muscle fibre number in utero. Biosci. Rep. 13, 213–220.

Remels, A.H.V., Langen, R.C.J., Schrauwen, P., Schaart, G., Schols, A.M.W.J., and Gosker, H.R. (2010). Regulation of mitochondrial biogenesis during myogenesis. Mol. Cell. Endocrinol. 315, 113–120.

Ritchie, M.E., Phipson, B., Wu, D., Hu, Y., Law, C.W., Shi, W., and Smyth, G.K. (2015). Limma powers differential expression analyses for RNA-sequencing and microarray studies. Nucleic Acids Res. 43, e47.

Robinson, M.D., and Oshlack, A. (2010). A scaling normalization method for differential expression analysis of RNA-seq data. Genome Biol. 11, R25.

Robinson, M.D., McCarthy, D.J., and Smyth, G.K. (2009). edgeR: A Bioconductor package for differential expression analysis of digital gene expression data. Bioinformatics 26, 139–140.

Shevchenko, A., Wilm, M., Vorm, O., and Mann, M. (1996). Mass spectrometric sequencing of proteins from silver-stained polyacrylamide gels. Anal. Chem. 68, 850–858.

Smyth, G.K. (2004). Linear Models and Empirical Bayes Methods for Assessing Differential Expression in Microarray Experiments. Stat. Appl. Genet. Mol. Biol. 3, 1–25.

Stockdale, F.E. (1992). Myogenic cell lineages. Dev. Biol. 154, 284–298.

Stöhr, G., and Tebbe, A. (2011). Chapter 8. Quantitative LC-MS of Proteins. pp. 104–122.

Tyanova, S., Temu, T., and Cox, J. (2016). The MaxQuant computational platform for mass spectrometry-based shotgun proteomics. Nat. Protoc. 11, 2301–2319.

Velloso, C.P. (2008). Regulation of muscle mass by growth hormone and IGF-I. Br.J.Pharmacol. 154, 557–568.

Wang, X., Jiang, L., Wallerman, O., Younis, S., Yu, Q., Klaesson, A., Tengholm, A., Welsh, N., and Andersson, L. (2018). ZBED6 negatively regulates insulin production, neuronal differentiation, and cell aggregation in MIN6 cells. FASEB J. fj.201600835R.

Wu, L., Timmers, C., Maiti, B., Saavedra, H.I., Sang, L., Chong, G.T., Nuckolls, F., Giangrande, P., Wright, F.A., Field, S.J., et al. (2001). The E2F1–3 transcription factors are essential for cellular proliferation. Nature 414, 457–462.

Yaffe, D., and Saxel, O. (1977). Serial passaging and differentiation of myogenic cells isolated from dystrophic mouse muscle. Nature 270, 725–727.

Younis, S., Kamel, W., Falkeborn, T., Wang, H., Yu, D., Daniels, R., Essand, M., Hinkula, J., Akusjärvi, G., and Andersson, L. (2018a). Multiple nuclear-replicating viruses require the stress-induced protein ZC3H11A for efficient growth. Proc. Natl. Acad. Sci. 115, 201722333.

Younis, S., Schönke, M., Massart, J., Hjortebjerg, R., Sundström, E., Gustafson, U., Björnholm, M., Krook, A., Frystyk, J., Zierath, J.R., et al. (2018b). The ZBED6-IGF2 axis has a major effect on growth of skeletal muscle and internal organs in placental mammals. Proc. Natl. Acad. Sci. U. S. A. 115, 201719278.

